# The miR-290 and miR-302 clusters are essential for reprogramming of fibroblasts to induced pluripotent stem cells

**DOI:** 10.1101/2024.09.02.610895

**Authors:** Julia Ye, Ryan M. Boileau, Ronald J. Parchem, Robert L. Judson-Torres, Robert Blelloch

**Author notes:** These authors contributed equally (Julia Ye, Ryan M. Boileau).

## Abstract

The miR-290 and miR-302 clusters of microRNAs are highly expressed in naïve and primed pluripotent stem cells, respectively. Ectopic expression of the embryonic stem cell-specific cell cycle regulating (ESCC) family of microRNAs arising from these two clusters dramatically enhances the reprogramming of both mouse and human somatic cells to induced pluripotency. Here, we used genetic knockouts to dissect the requirement for the miR-290 and miR-302 clusters during the reprogramming of mouse fibroblasts into induced pluripotent stem cells (iPSCs) with retrovirally introduced Oct4, Sox2, and Klf4. Knockout of either cluster alone did not negatively impact the efficiency of reprogramming. Resulting cells appeared identical to their embryonic stem cell microRNA cluster knockout counterparts. In contrast, the combined loss of both clusters blocked the formation of iPSCs. While rare double knockout clones could be isolated, they showed a dramatically reduced proliferation rate, a persistent inability to fully silence the exogenously introduced pluripotency factors, and a transcriptome distinct from individual miR-290 or miR-302 mutant ESC and iPSCs. Taken together, our data show that miR-290 and miR-302 are essential yet interchangeable in reprogramming to the induced pluripotent state.

**Impact Statement:** The process by which somatic cell reprogramming yields induced pluripotent stem cells (iPSCs) is incompletely understood. MicroRNAs from the miR-290 and miR-302 clusters have been shown to greatly increase reprogramming efficiency, but their requirement in the process has not been studied. Here, we examine this requirement by genetically removing the miRNA clusters in somatic cells. We discover that somatic cells lacking either, but not both, of these miRNA clusters can form iPSC cells. This work thus provides new important insight into mechanisms underlying reprogramming to pluripotency.

## Introduction

The derivation of induced pluripotent stem cells (iPSCs) from somatic cells through the introduction of exogenous factors^1^ has been a major boon to the scientific and medical communities, as it has enabled new and previously unimaginable advancements in disease modeling, pharmaceutical development, and regenerative medicine (reviewed in ^2–6^). However, the complex interactions among the molecular events that drive somatic cell reprogramming or de-differentiation to iPSCs remain incompletely understood (reviewed in ^7,8^). MicroRNAs (miRNAs), which are small non-coding RNAs that destabilize and inhibit the translation of mRNAs^9–11^, have been found to play prominent roles in the reprogramming process.

The *mir-290~295* (*mir-371~373* in humans^12^) and *mir-302~367* clusters of miRNAs, in particular, have been studied extensively in pluripotent cells, including iPSCs. Along with *mir-17~92* and *mir-106b~25*, they are the most highly expressed miRNA clusters in pluripotent stem cells (reviewed in ^13–15^). Specifically, miR-290~295 (miR-290) miRNAs are highly expressed in “naïve” pluripotent mouse embryonic stem cells (ESCs) and primordial germ cells^16–18^, while miR-302~367 (miR-302) miRNAs are highly expressed in “primed” pluripotent mouse epiblast stem cells (mEpiSCs) and human ESCs^19–21^. Many of the individual miRNAs in these clusters share a similar seed sequence (AAGUGC) and thus belong to the same miRNA family—the embryonic stem cell-specific cycle regulating (ESCC) family of miRNAs^22,23^. The common seed sequence also suggests common mRNA targets among these miRNAs^24^.

The miR-290 and miR-302 miRNAs are intimately integrated within the pluripotency network. Their promoters are bound by the core pluripotency transcription factors, Oct4, Sox2, Nanog, and Tcf3^17,25^ Among their many functions in pluripotent cells (reviewed in ^15^), these miRNAs suppress the G1-S checkpoint^22^, inhibit the silencing of ESC self-renewal^26,27^, and enable the transition between alternative pluripotent states^28^. Notably, introducing the ESCC miRNAs dramatically enhances reprogramming by the Yamanaka factors in both mouse and human somatic cells^29,30^. Indeed, it has been shown that overexpression of miR-302 cluster alone is capable of inducing mouse and human reprogramming^31,32^. The *mir-290~295* and *mir-302~367* cluster-derived miRNAs promote iPSC generation through a number of different mechanisms and downstream targets, including cell signaling, the mesenchymal-to-epithelial transition, cell cycle, epigenetic modifiers, endoplasmic reticulum trafficking, and cellular metabolism^33–36^.

Given the remarkable impact of exogenous ESCCs on reprogramming and their high levels of expression in pluripotent stem cells, we wanted to ask whether they are necessary for somatic cell reprogramming. To do so, we used genetic knockouts (KO) for the *mir-290~295* (*mir-290* KO^37^) and *mir-302a-d* (*mir-302* KO^38^) loci, as well as a CRISPR-mediated deletion of the entire *mir-302~367* locus. Surprisingly, deletion of either miRNA locus alone did not diminish the formation of iPSC-like colonies. The resulting cells expressed endogenous pluripotency markers at similar levels as wild-type PSCs and silenced the retrovirally introduced reprogramming factors, consistent with complete reprogramming. However, given the incomplete developmental potential of *mir-290* and *mir-302* KO PSCs^37–41^, we refer to these cells as iPSC-like. It was also possible to generate and culture cells from fibroblasts deficient for both the *mir-290~295* and *mir-302~367* loci. However, the resulting double KO cells show signs of incomplete maturation, including proliferation defects, an inability to fully silence the exogenously introduced reprogramming factors, and a transcriptomic profile distinct from that of ESC and iPSC-like cells deficient in either miR-290 or miR-302. Based on these findings, we propose that *mir-290~295* and *mir-302~367* clusters are individually dispensable but function with redundancy in their control of the maturation of the induced pluripotent state.

## Results

### miR-302 is not required for reprogramming

To investigate whether miR-302 miRNAs are required for the de-differentiation of somatic cells to iPSCs, we harvested mouse embryonic fibroblasts (MEFs) from *mir-302* KO embryos. In this KO model, eGFP was knocked into the *mir-302~367* locus, such that *mir-302a-d* are deleted, while *mir-367* is left intact (Figure 1A^38^). MEFs were transduced with retroviruses expressing the pluripotency factors Oct4, Sox2, Klf4 (OSK) and cultured in ESC media (Figure 1B). Resulting cultures were stained 15 days post-transduction for Nanog, a marker of cells reprogrammed to the iPSC state. We found that *mir-302* KO MEFs were able to generate Nanog+ colonies at efficiencies similar to their control *mir-302* heterozygous (Het) counterparts (Figure 1C and D). The dispensability of endogenous miR-302 was surprising given previous work showing a requirement for the locus in the production of human induced pluripotent stem cells^42^; however, it is consistent with other work showing that mouse iPSC clones can form without activation of the endogenous miR-302 locus^18^.

**Figure 1.**
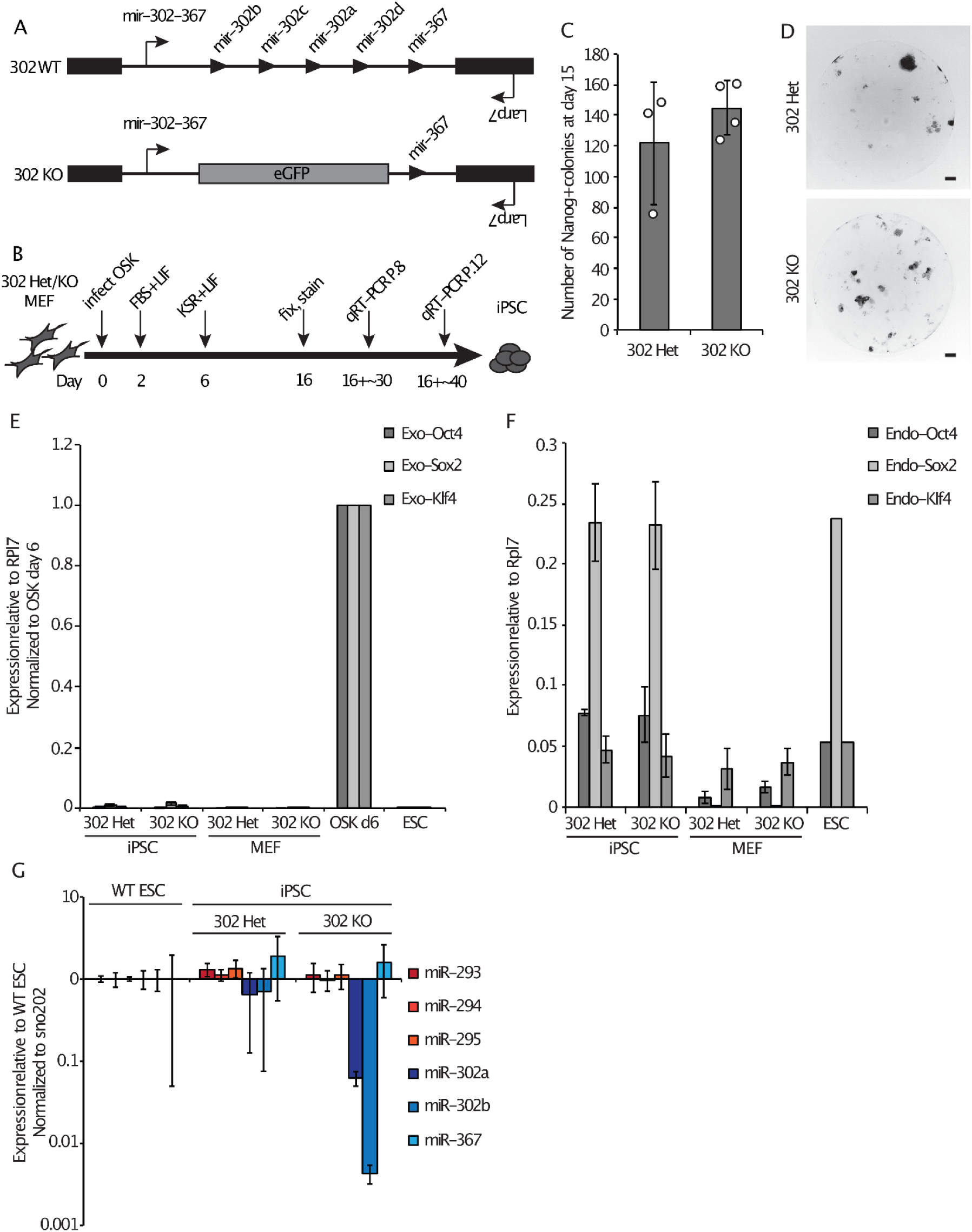
miR-302 is not required for reprogramming. (A) Schematic of the WT and KO alleles of the *mir-302~367* locus as previously described^38^. Note that *mir-367* is not affected in this *mir-302* KO design. (B) Schematic of the reprogramming protocol and iPSC analysis time points. (C) Average number of Nanog+ colonies counted per well 15 days after OSK transduction. (D) Representative images of individual Nanog-stained OSK reprogramming wells at day 15 taken at 2x magnification (scale bar represents 500 um). (E) qRT-PCR analysis of exogenous OSK expression levels in WT ESCs, and *mir-302* Het and *mir-302* KO MEFs and iPSC-like cells analyzed ~30 days (P.8) after reprogramming day 16. Data are normalized to MEFs collected 6 days after OSK transduction. (F) qRT-PCR analysis of endogenous OSK expression levels in WT ESCs and *mir-302* Het and *mir-302* KO MEFs and P.8 iPSC-like cells. (G) Transcript levels of mature *mir-290~295* and *mir-302~367* cluster miRNAs in WT ESCs, *mir-302* Het iPSCs, and *mir-302* KO iPSC-like cells at P.12. Error bars represent SD of 3-4 biological replicates. Circles indicate individual data points.

We expanded iPSC-like cells arising from the control and *mir-302* KO MEFs and verified the genetic status of each line (Supplemental Figure 1A). After ~30 days in culture (passage 8, P.8), we analyzed iPSC gene expression and saw that both *mir-302* Het and *mir-302* KO cells appropriately silenced exogenous retrovirus expression (Figure 1E) and activated endogenous pluripotency genes (Figure 1F). They are indistinguishable from ESCs morphologically (Supplemental Figure 1B) and have high colony formation efficiencies expected of self-renewing pluripotent cells (Supplemental Figure 1C). Additionally, *mir-302* Het and KO cells express similar levels of the *mir-290~295* cluster miRNAs compared to wild-type (WT) ESCs, demonstrating that there is no compensatory increase in miR-290 levels in the absence of miR-302 (Figure 1G). Expression of a representative panel of naïve and primed pluripotency markers is similar across WT ESCs, *mir-302* Het cells, and *mir-302* KO cells, except for a trend (not statistically significant) toward increased Rex1 levels in Hets vs. KO (Supplemental Figure 1D). Together, these data show that miR-302 is dispensable for both achieving and maintaining an induced pluripotent-like state.

### Loss of miR-290 leads to slightly delayed iPSC maturation and a compensatory increase in miR-302

Next we asked whether the miR-290 miRNAs are required for reprogramming to iPSCs. Unlike miR-302, miR-290 is highly expressed in naïve pluripotent stem cells, the endpoint of reprogramming^16–18^. MEFs were isolated from embryos in which the entire *mir-290~295* locus had been replaced with a pPGK-Neo cassette (*mir-290* KO, Figure 2A^37^). Control *mir-290* Het and *mir-290* KO MEFs were transduced with OSK and then stained 15 days later for Nanog as described above (Figure 2B). The *mir-290* KO cells showed no significant difference in the number of Nanog+ colonies formed compared with control (Figure 2C and 2D). Resulting colonies were expanded and their genotypes were confirmed by PCR (Supplemental Figure 2A).

**Figure 2.**
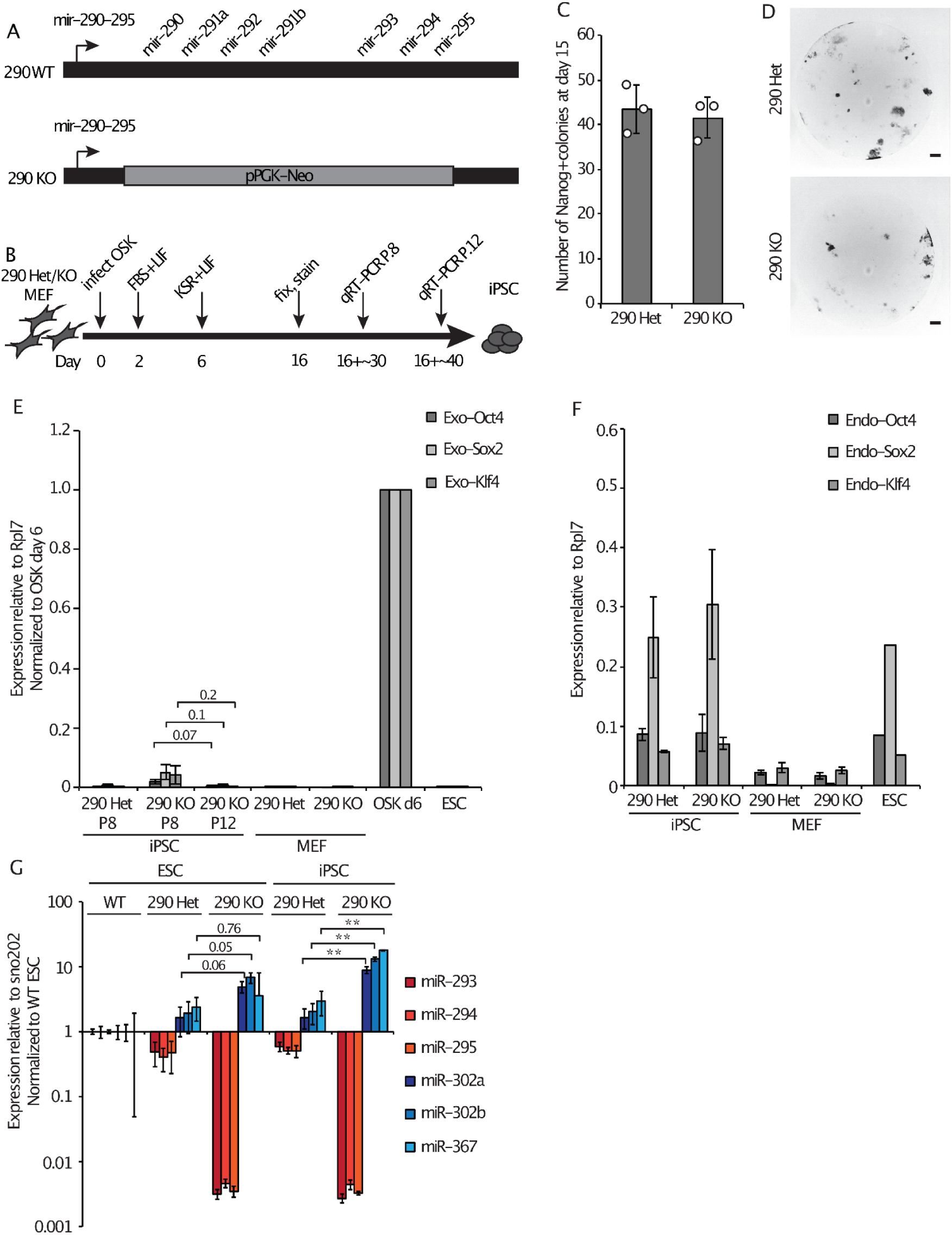
Loss of miR-290 leads to slightly delayed iPSC maturation and a compensatory increase in miR-302. (A) Schematic of the WT and KO alleles of the *mir-290~295* locus as previously described^37^. (B) Schematic of the reprogramming protocol and iPSC analysis points. (C) Average number of Nanog+ colonies counted per well 15 days after OSK transduction. (D) Representative images of individual Nanog-stained OSK reprogramming wells at day 15 taken at 2x magnification (scale bar represents 500 um). (E) qRT-PCR analysis of exogenous OSK expression levels in WT ESCs, and *mir-290* Het and *mir-290* KO MEFs and iPSC-like cells at P.8 or P.12. Data are normalized to MEFs collected 6 days after OSK transduction. (F) qRT-PCR analysis of endogenous OSK expression levels in WT ESCs and *mir-290* Het and *mir-290* KO iPSC-like cells at P.8. (G) Transcript levels relative to internal control and normalized to WT ESCs of mature *mir-290~295* and *mir-302~367* cluster miRNAs in WT ESCs, *mir-290* Het ESCs or P.12 iPSC-like cells, and *mir-290* KO ESCs or P.12 iPSC-like cells. Error bars represent SD of 3 biological replicates. (**) P < 0.001; numbers above the graphs in E and G indicate the P-values of the comparisons marked, two-sided t-test. Circles indicate individual data points.

We observed that the morphology of the *mir-290* KO colonies was slightly abnormal (Supplemental Figure 2B). Specifically, cells within the colonies were morphologically more heterogeneous, with a mix of compact cells resembling WT ESCs, as well as spread out cells growing out from colony borders. To determine whether this phenotype was due to a defect in the reprogramming process or intrinsic to *mir-290* deficient pluripotent stem cells, we derived embryonic stem cells (ESCs) from *mir-290* KO blastocysts. The resulting *mir-290* KO ESCs, but not WT or Het controls, showed a similar slightly abnormal morphology as the *mir-290* KO iPSC-like cells (Supplemental Figure 2B). A colony reforming assay also showed a reduced efficiency in colony formation both in *mir-290* KO iPSC-like cells and in *mir-290* KO ESCs (Supplemental Figure 2C). This diminished self-renewal capacity likely represents a propensity for spontaneous differentiation, consistent with the flattened morphology of some of the cells.

Expanded *mir-290* Het and KO iPSC-like cells silenced the exogenously introduced pluripotency factors, while activating the endogenous pluripotency genes (Figure 2E, 2F, Supplemental Figure 2D), consistent with maturation of the iPSC clones^43–46^. The *mir-290* KO cells showed a slight delay in the silencing of the exogenous factors, as there was ongoing low-level expression at P.8, which subsequently decreased to undetectable levels 13-15 days later at P.12 (Figure 2E). miR-290-deficient ESCs and iPSCs showed a dose-dependent upregulation of miR-302, with increasingly elevated miR-302 levels observed with loss of each *mir-290* allele (Figure 2G). Together, these data show that miR-290, like miR-302, is not alone necessary for reprogramming to an iPSC-like state.

### miR-290 and miR-302 double knockout cells fail to reprogram to iPSCs

Given the upregulation of miR-302 in *mir-290* KO cells and the shared seed sequence among most of the miRNAs in the two clusters, we next asked whether these two loci function redundantly duringreprogramming. Because *mir-290;mir-302* double KO embryos arrest during early post-implantation development, it is impossible to isolate MEFs from these animals^38^. Furthermore, attempts at using a miR-302 sponge failed to fully block miR-302 function as determined by measurement of known targets (data not shown). Therefore, we devised a CRISPR-based strategy to genetically remove miR-302 in *mir-290* KO fibroblasts. Guide RNAs (gRNAs) were designed to flank the 5’ and 3’ ends of just *mir-302a-d* (gRNAs g1 and g2) or the entire *mir-302~367* locus (gRNAs g1 and g3) to also test the impact of *mir-367* loss on the reprogramming phenotype (Figure 3A). MEFs were harvested from *mir-290* WT, Het, and KO embryos and electroporated with the CRISPR constructs to induce *mir-302a-d* or *mir-302~367* deletions (Figure 3B). As a control, we also electroporated CRISPR constructs containing no gRNAs (“empty”). The CRISPR constructs contain GFP and BFP markers, which were used to sort for MEFs that received both guide RNAs. Given the limited number of population doublings that MEFs can undergo before they senesce, it was not possible to subclone the CRISPR transfected MEFs. However, DNA genotyping of the sorted bulk MEF populations showed efficient deletion of the *mir-302a-d* and *mir-302~367* loci (Supplemental Figure 3A).

**Figure 3.**
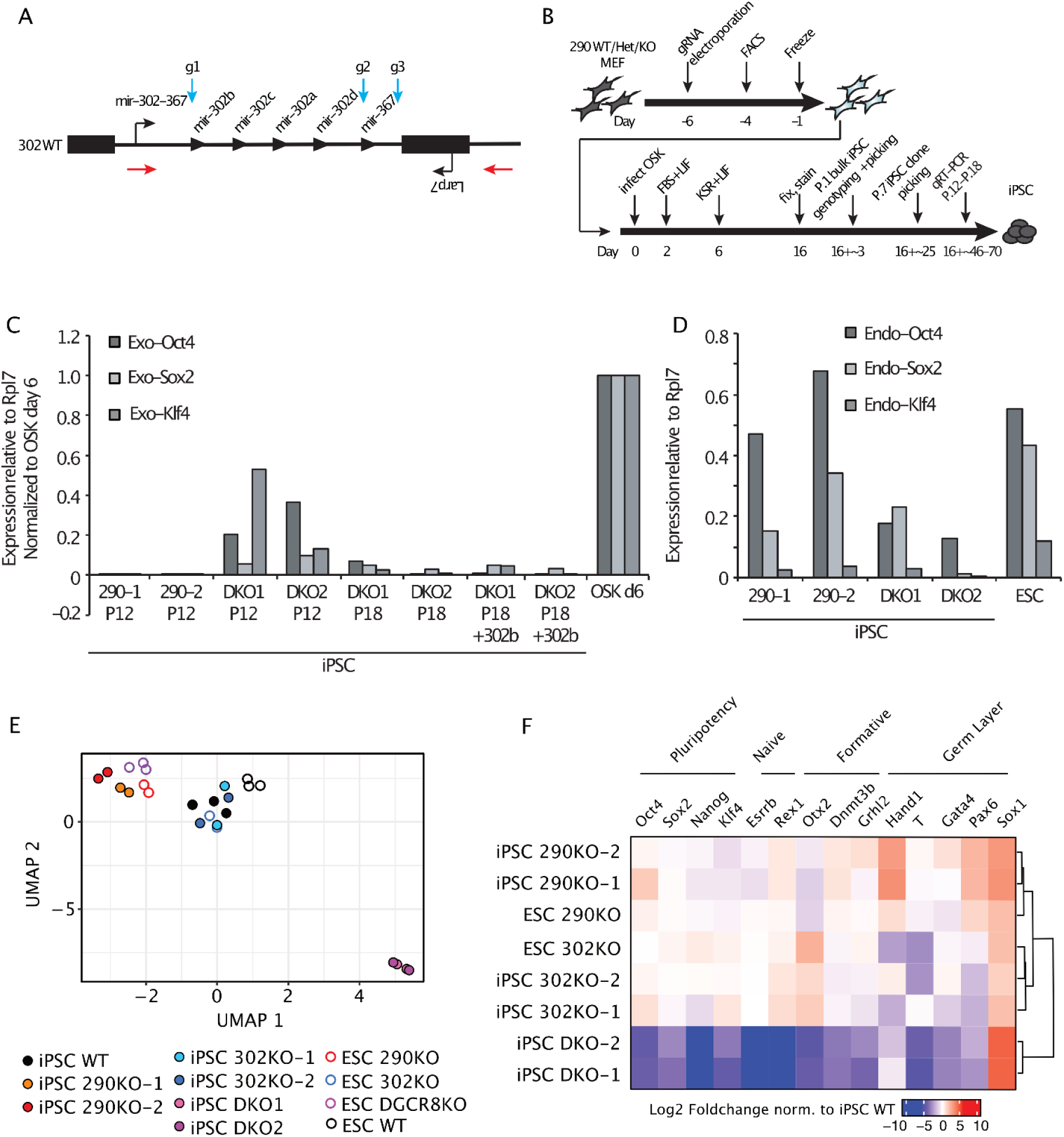
miR-290 and miR-302 double knockout cells fail to reprogram to iPSCs. (A) Schematic of the CRISPR deletion strategy for the *mir-302~367* locus. Blue arrows indicate the location of gRNAs g1, g2, and g3. Red arrows indicate the location of the PCR primers used for DNA genotyping. (B) Schematic of the reprogramming protocol and iPSC analysis time points. (C) qRT-PCR analysis of exogenous OSK expression levels in *mir-290−/−;mir-302~367+/+* (290-1, 290-2) and *miR-290−/−;mir-302~367−/−* (DKO1, DKO2) iPSC-like cell lines at P.12 and at P.18 in the presence or absence of exogenous miR-302b. Data are normalized to MEFs collected 6 days after OSK transduction. (D) qRT-PCR analysis of endogenous OSK expression levels in *mir-290−/−;mir-302~367+/+* (290-1, 290-2) and *miR-290−/−;mir-302~367−/−* (DKO1, DKO2) iPSC lines at P.12. (E) UMAP clustering analysis of RNA-seq data for ESCs and iPSC-like cell lines at P.18. (F) Heatmap of gene expression for select markers of pluripotent cell identity. Data is shown as Log2 Foldchange relative to iPSC WT and is hierarchically clustered by sample.

We transduced the FACS-enriched gRNA-treated MEFs with OSK retroviruses as before and stained for Nanog expression 16 days later (Figure 3B). Deletion of *mir-302a-d* in a *mir-290* WT background had no impact on the number of Nanog+ colonies formed (Supplemental Figure 3B), which mirrors our results using MEFs derived from *mir-302a-d* knockout animals (Figure 1C). Extending these results, the deletion of the entire cluster including *mir-367* also did not alter the number of Nanog+ colonies. Similar results were seen in the *mir-290* Het background. Surprisingly, the deletion of *mir-302a-d* or *mir-302~367* in the *mir-290* KO background also did not reduce the number of Nanog+ colonies formed compared to WT. These results suggest that even in the absence of all ESCC miRNAs, fibroblasts can still efficiently form Nanog-positive colonies.

Next, we asked if these Nanog-positive colonies are stable and produce mature iPSC lines as defined by their ability to be passaged and silence the exogenously introduced pluripotency factors. Genotyping of the bulk population at P.1 showed very similar ratios of *mir-302~367* WT to KO alleles as in the originating MEFs, supporting the interpretation that deleting the two clusters does not result in a competitive disadvantage of the cells in producing the Nanog-positive colonies (Supplemental Figure 3C, top panel). However, *mir-302a-d* KO and *mir-302~367* KO alleles were lost in the *mir-290* KO background after six more passages (Supplemental Figure 3C, bottom panel), while all other genotypes maintained similar ratios as seen at P.1. To further characterize this finding, clones were picked for each condition at P.7, expanded, and genotyped (Supplemental Figure 3D). Consistent with bulk population data, it was possible to isolate all possible genotypes when starting with the *mir-290* WT and *mir-290* Het cells. In contrast, we were able to isolate only one *mir-290−/−;mir-302~367+/-* colony, and we were unable to identify any *mir-290* KO clones with concurrent *mir-302a-d* or *mir-302~367* deletion, despite screening dozens of additional colonies to search for rare clones.

The absence of double knockout at P.7 could result from unedited mir-290KO cells outcompeting double KO cells over time in culture. To address this possibility, we picked individual colonies at P.1 immediately after the conclusion of the reprogramming assay. We were able to identify at least 2 clones of *mir-290;mir302~367* double KO cells as confirmed by DNA genotyping and miRNA expression (Supplemental Figure 3E and 3F), thus demonstrating that miR-290 and miR-302 are not absolutely required for the generation and maintenance of cell lines. However, these cell lines appeared distinct from the iPSC-like lines recovered from single KOs, showing a more flattened and dispersed morphology (Supplemental Figure 3G). Crystal violet proliferation assays showed that the double KO cells had a severe proliferation defect relative to *mir-290* single knockout cells (Supplemental Figure 3H). Mixing equal numbers of *mir-290;mir302~367* double KO cells and *mir-290* single KO iPSCs led to the rapid loss of double KO genotypes, similar to what we had seen in bulk iPSC populations (Supplemental Figure 3E). qRT-PCR analysis of the double KO cell lines revealed impaired silencing of the retrovirally expressed pluripotency factors, which was mitigated but not fully corrected after extended culture and passaging to P.18 (Figure 3C). This phenomenon was observed even upon transient reintroduction of the ESCC miR-302b for 8 days (Figure 3C). Moreover, the double KO iPSCs showed reduced expression of the endogenous pluripotency factors despite extended passaging (Figure 3D).To further assess the reprogramming status of the double and single miR-290 and miR-302 KO cell lines, we performed RNA-seq. To differentiate expression changes associated with reprogramming defects versus the direct impact of the miRNAs, we included *mir-290* KO and *mir-302* KO ESCs as controls. We also incorporated RNA-seq data from *DGCR8* KO ESCs which lack all mature miRNAs^47^, as the closest control we had to the double KO lines. Clustering using UMAP analysis revealed several important findings (Figure 3E). First, *mir-290* KO iPSC-like cells clustered with *mir-290* KO ESCs, consistent with complete reprogramming to an ESC-like state. *mir-290* KO pluripotent cells (both ESC and iPSC lines) clustered separately from their corresponding WT cells, which is expected given that the miRNAs from this cluster make up the majority of miRNAs in ESCs, with well-known effects on many downstream transcripts^13–15^. In contrast, *mir-302* KO ESCs and iPSC-like cells clustered with WT ESCs, consistent with full reprogramming in the absence of miR-302. Unlike *miR-290* KO, the clustering of *miR-302* KO iPSCs with WT ESCs is expected of fully reprogrammed cells given that this cluster is lowly expressed in ESCs. ESCs lacking *DGRC8* clustered most closely with *mir-290* KO cells, consistent with the predominance of miRNAs in ESCs coming from the miR-290 cluster. However, double KO cell lines demonstrated a transcriptomic profile that was distinct from that of all other genotypes, consistent with a failure in the completion of reprogramming.

Differential gene expression analysis uncovered ~6000 differentially expressed genes in the double KO cells compared to WT iPSCs (Log2 Foldchange > 1 and adjusted p.value < 0.05)(Supplemental Figure 4A). This was dramatically more than those found for *mir-290*KO iPSC-like cells (~1000 genes) and for *mir-302*KO lines (43 genes). Clusterprofiler analyses of gene ontologies for differentially expressed genes in *mir-290*KO iPSC-like cells suggested abnormal upregulation of genes involved in neural identity (Supplemental Figure 4B). Similar categories of genes were enriched in double KO iPSC-like cells. However, in addition, there was an enrichment for genes related to mesoderm development (Supplemental Figure 4C), consistent with the mesoderm origin of MEFs. No categories were enriched in *mir-302*KO lines. We also evaluated the expression of predicted *mir-290/302* family targets in our iPSC-like mutants (Supplemental Figure 5A). Relative to all other genes, predicted *mir-290/302* family targets were elevated in 290KO ESC/iPSC mutant lines and greatly elevated in double KO iPSCs. We did not observe similar increases in *mir-302*KO ESCs or iPSC-like cells. We next evaluated the expression of specific genes related to pluripotent identity, including naïve, formative, and germ layer markers (Figure 3F and Supplemental Figure 5B). Pluripotency genes Oct4, Sox2, Nanog, and Klf4 were highly expressed in all *mir-290*KO or *mir-302*KO cell lines. However, *mir-290*KO ESC and iPSC-like lines all had higher levels of the germ layer markers Hand1, Pax6, and Sox1 relative to *mir-302*KO and WT iPSCs. In the double KO cells, Oct4, Sox2, Nanog, and Klf4 were lowly expressed relative to all other cell lines. Further, we found most markers of pluripotent states were not expressed, with a notable exception of Sox1 which was highly upregulated relative to other lines. To directly test their developmental potential, we attempted to make embryoid bodies (EBs). However, EBs from the double KO cells degenerated by day 4 before germ layer markers would be expected to be expressed. Single knockouts formed EBs but were morphologically distinct from WT given their known developmental phenotypes (data not shown)^37,38^. Together, these data uncover an absolute requirement for either miR-290 or miR-302 in reprogramming to a mature iPSC state.

## Discussion

In this study, we used genetic KO models of the miR-290 and miR-302 miRNA families to test their requirement in the reprogramming of mouse somatic cells to iPSCs. Given the remarkable impact of exogenously introduced microRNAs from these clusters on the reprogramming of fibroblasts to iPSCs^29,30,33^, it was surprising to discover that the absence of either miR-290 or miR-302 had little effect on the formation of iPSC-like colonies. Nanog+ colonies formed in numbers and with kinetics similar to that seen with their control counterparts; resulting cells self-renewed stably and proliferated at WT rates. Furthermore, the retrovirally introduced exogenous factors were ultimately silenced, an important demonstration of iPSC maturation^43–46^. Consistent with complete maturation, gene expression showed them to be identical to their ESC counterparts. These results show that neither cluster alone is required for the reprogramming of mouse fibroblasts to iPSC-like cells.

We describe these cells as “iPSC-like,” because, given the known roles for miR-302 and miR-290 in embryonic development, we would not expect them to have full developmental competence. For example, *mir-302~367* locus is required for proper development of various somatic organs^38,40^, and miR-290 plays important roles in placental and germline development^37,38,48^. Furthermore, the absence of miR-290, whether in the resulting iPSCs or independently in ESCs derived from knockout epiblasts, showed a metastable like state where cells showed mixed morphologies including both dome like colonies consistent with the naïve pluripotent state as well as flatter cells more consistent with some early differentiation^49^. Indeed, bulk sequencing of the miR-290 KO cells showed elevated levels of some early differentiation markers such as Hand1, Pax6, and Sox1 while also expressing high levels of pluripotency markers. In contrast to the single knockouts, deletion of both clusters failed to produce iPSC-like cells. While they produced Nanog+ colonies, expansion of double knockout cells showed defects in proliferation, silencing of retrovirally expression, and a highly abnormal expression profile including reduced expression of pluripotency markers and elevated expression of mesoderm markers found in the originating fibroblast population. Therefore, the two clusters are redundant in terms of reprogramming; that is, while loss of either cluster alone is compatible with full reprogramming to an iPSC state, the loss of both is not. Given that the two clusters share the expression of the ESCC family of microRNAS, it is likely these miRNAs drive the requirement for either cluster in reprogramming. However, complex combinatorial knockouts of all the ESCC miRNAs without disruption of non-ESCC miRNAs in the two clusters will be required to formally prove this assumption.

**Figure 4:**
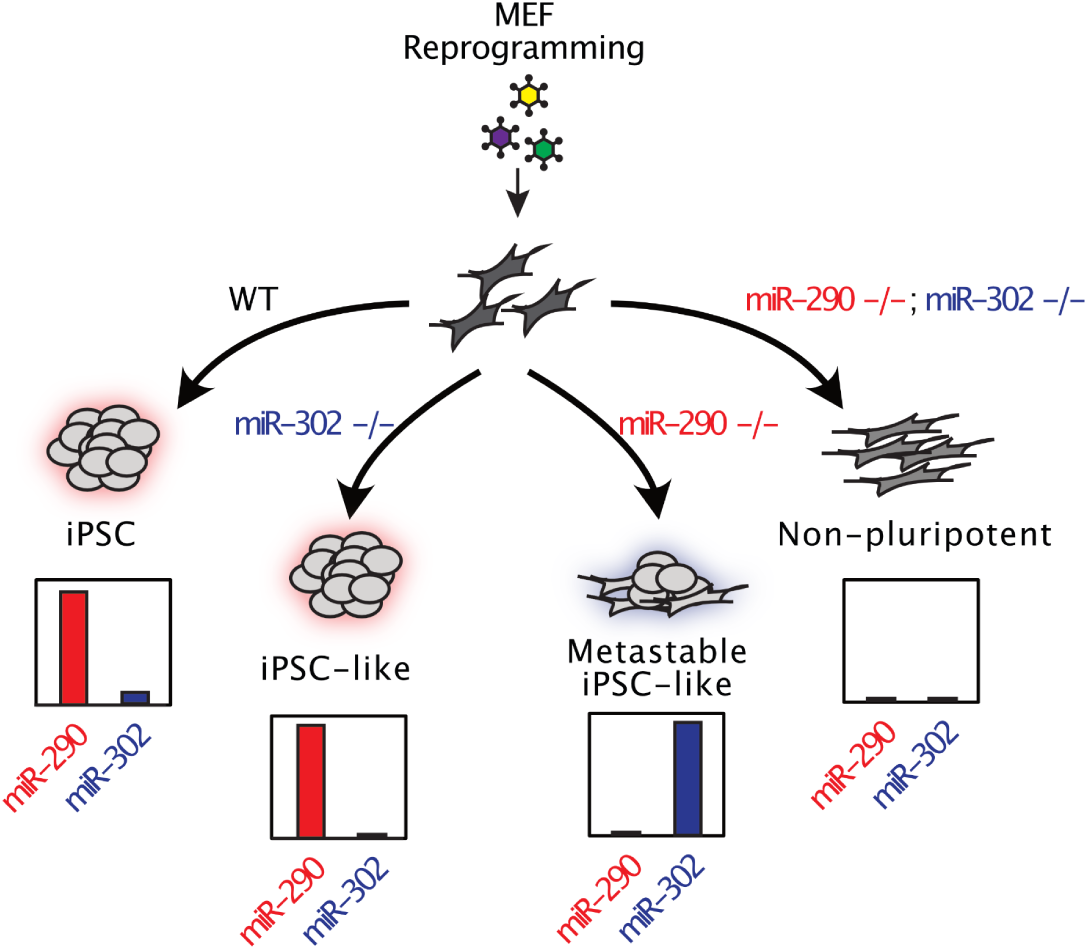
Graphical Summary. The miR-290 and miR-302 clusters are individually dispensable but combined essential for the reprogramming of somatic cells to an induced pluripotent state. OSK-reprogrammed miR-302 KO fibroblasts appear identical morphologically and expression-wise to WT iPSCs/ESCs. OSK-reprogrammed miR-290 KO fibroblasts show increased morphological heterogeneity and premature expression of some early differentiation markers including miR-302 relative to WT iPSCs/ESCs, but are identical to their miR-290 KO ESC counterparts, showing they can reach a mature iPSC-like state. In contrast, miR-290;miR-302 double knockout fibroblasts cannot be reprogrammed to a mature pluripotent-like state.

Our findings contrast with other studies showing an essential role for the miR-302 cluster in the reprogramming of human fibroblasts to iPSCs^42^. This difference may reflect the distinct final states of mouse and human iPSCs. Specifically, mouse iPSCs represent a naïve pluripotent state^50,51^, which is characterized by high levels of miR-290 but not miR-302~367^16–18^. In contrast, human ESCs are more similar to mouse EpiSCs, which are grown in FGF and activin and are considered a primed pluripotent state^52,53^. Primed pluripotent cells express miR-302~367 and significantly reduced levels of miR-371, the human miR-290 ortholog^19–21^; therefore, it is possible that the *mir-302~367* cluster would be required to produce induced mEpiSCs^54^. Going forward, it will be interesting to dissect the impact of miR-302 loss on reprogramming mouse fibroblasts to an induced EpiSC state.

Prior work using microRNA biogenesis mutants have produced conflicting results. Conditional deletion of Dicer was shown to block transcription factor-induced reprogramming of MEFs to iPSCs^54,55^. In contrast, conditional deletion of *Dgcr8* was shown to suppress but not fully block transcription factor induced reprogramming of MEFs^56^. One possible explanation for this discrepancy is the known role for Dicer in the biogenesis of other short RNA species in ESCs including shRNAs and mirtrons^56,57^. However, our findings showing that the loss of both miR-290 and miR-302 blocks reprogramming supports a requirement of miRNAs in the induction of pluripotency. A likely explanation for the reported results in *Dgcr8* deletion study is that any delay in the timing of deletion of the gene and loss of protein could allow for rare cells that are still producing enough miRNAs including those derived from either miR-302 and miR-290 clusters to become iPSCs. Indeed, given the requirement for miRNAs in the survival of MEFs, the biogenesis experiments required a balance between keeping the source cells healthy enough while depleting as many miRNAs as possible upon the induction of reprogramming. Given that miR-290 and miR-302 are not expressed in MEFs, this was not an issue in our study. Therefore, the combined results of these studies support a transient requirement for miR-290 and miR-302 during reprogramming. Once in a pluripotent state, stem cells do not require the biogenesis enzymes to maintain their self-renewal, although they are required for their normal differentiation as are the miR-290 and miR-302 clusters^58–60^.

## Conclusion

In sum, our work uncovers an absolute requirement for the miR-290 and miR-302 clusters in iPSC reprogramming. Either cluster alone is enough to enable the production of iPSCs with little effect on efficiency and self-renewal of the final iPSC clones. These iPSC clones are essentially identical to their ESC counterparts. The miRNAs from these clusters have been implicated in a host of critical activities in pluripotent stem cells and are powerful promoters of de-differentiation^26–33,36^. Understanding the targets of these miRNAs underlying their requirement will continue to provide important mechanistic insight into cellular events essential to the reprogramming to pluripotency^33,36^.

## Materials and Methods

### MEF generation

MEFs were generated as previously described^29^. In brief, WT, *mir-302+/-*, *mir-302−/−*, *mir-290+/-*, and *mir-290−/−* embryos were harvested at E13.5. After removal of the head and visceral tissue, the remaining tissue was dissociated by physical disruption and trypsinization and plated as Passage 0 (P0) cells in MEF medium (high glucose (H-21) DMEM, 10% FBS, non-essential amino acids, L-glutamine, penicillin/streptomycin, and 55uM beta-mercaptoethanol). MEFs were expanded to P3 and frozen.

For generation of *mir-290/mir-302* double KO MEFs, CRISPR^61^ was used to delete *mir-302a-d* or *mir-302~367* in *mir-290+/+*, *mir-290+/-*, and *mir-290−/−* MEFs. In brief, guide RNAs (sequences designed using crispr.mit.edu) were inserted into the PX458 construct or a modified PX458 in which eGFP was replaced by BFP. Simultaneous introduction of gRNAs g1 (CACC-G-TTAACTAGTTGCCTTGTGGG, AAAC-CCCACAAGGCAACTAGTTAA-C) and g2 (CACC-GGAGCCACCACACTCAAACA, AAAC-TGTTTGAGTGTGGTGGCTCC) was used to create the *mir-302a-d* KO allele. Simultaneous introduction of gRNAS g1 and g3 (CACC-G-TTGCACTTTAGCAATGGTGA, AAAC-TCACCATTGCTAAAGTGCAA-C) was used to create the *mir-302~367* KO allele. P0 *mir-290+/+*, *mir-290+/-*, and *mir-290−/−* MEFs were electroporated with the g1+g2 or g1+g3 constructs (~7.5ug of each construct per 3 million cells) using the Neon Transfection System (Invitrogen) according to the manufacturer’s protocol (1350V, 30ms, 1 pulse). Electroporated MEFs were sorted for GFP+BFP+ cells by fluorescence-activated cell sorting (FACS) 2 days later (BD FACS Aria3u), expanded, and frozen 3 days after FACS.

### Virus production

For retrovirus production, HEK293T cells grown in MEF medium were seeded at 2 million cells/10cm plate and transfected the following day with 5ug pCL-Eco and 5ug pMXs-expression plasmid with 30ul Fugene 6. Medium was replaced 24hr after transfection, and at 48hrs post-transfection, supernatant was harvested, filtered (0.45um), aliquoted, and frozen at −80C. Fresh aliquots were used for each experiment.

### Reprogramming/de-differentiation

P4 MEFs were plated onto 0.2% gelatin-coated Greiner uClear black-walled 96-well imaging plates at 2000 cells/well or standard 12-well plates at 20,000 cells/well. The next day (day 0), cells were treated with 40ul (96-well) or 400ul (12-well) of each retrovirus aliquot and 4ug/ml polybrene. At day 1, the medium was replaced with fresh MEF medium, and medium was replenished every other day with FBS+LIF medium (DMEM, 15% FBS, non-essential amino acids, L-glutamine, penicillin/streptomycin, 55uM beta-mercaptoethanol, 1000U/ml LIF) between days 2 and 6. Thereafter, medium was replenished with KSR+LIF medium (Knock-out DMEM (Invitrogen), 15% Knock-out serum replacement (Invitrogen), non-essential amino acids, L-glutamine, penicillin/streptomycin, 55uM beta-mercaptoethanol, 1000U/ml LIF). Reprogramming plates were analyzed for successful iPSC generation at day 15 or 16 as by defined by positive Nanog staining (CST8785). Immunostained wells were imaged using the INCell Analyzer 2000 (GE), and Nanog+ colonies were counted.

### ESC and iPSC culture and analyses

ESC lines (*mir-290+/+*, *mir-290+/-*, and *mir-290−/−*) were derived from individual E3.5 blastocysts plated in one well of a 24-well dish on a MEF feeder layer using ES medium with 20% KSR and 1 μM PD0325901. Blastocyst outgrowths were trypsinized and passaged after 3-4 days until ESC lines were established. iPSC-like cell lines were generated from reprogramming experiments described above.

ESC and iPSC-like cell lines were expanded and passaged under feeder-free conditions on 0.2% gelatin-coated plates in FBS+LIF medium (DMEM, 15% FBS, non-essential amino acids, L-glutamine, penicillin/streptomycin, 55uM beta-mercaptoethanol, 1000U/ml LIF) and depleted of un- or partially-reprogrammed MEFs by transferring un-adhered cells to new plates 30min-1hr after passage.

ESC and iPSC colony formation/clonogenicity assays were performed after the reprogramming protocol (P.18+). After adapting cells to FBS+LIF+2i (FBS+LIF and 1uM PD0325901, 3uM CHIR99021) we plated cells in 96-well plates at 500 cells/well and stained with Nanog antibody (CST8785) to count Nanog+ colonies. The ratio of Nanog+ colonies formed to the number of cells plated was defined as the colony formation potential.

To isolate and genotype individual iPSC-like clones for the CRISPR guide RNA-treated cells, iPSC-like cells were plated at 10,000 cells (for *mir-290+/+* and *mir-290+/-* cells) or 30,000 cells (for *mir-290−/−* cells) per 10cm plate and grown for 6-7 days in the feeder-free conditions described above. Individual colonies were physically picked and placed into 96 well plates, expanded, and harvested for DNA genotyping.

For mixing experiments, 100,000 of each cell line (*mir-290−/−;mir-302~367+/+* or *mir-290−/−;mir-302~367−/−*) were combined for 200,000 total cells per well of a 6-well plate in FBS+LIF medium. Every 3 days, cells were passaged at 200,000 cells per well, and the remaining cells were harvested for DNA genotyping analysis. The genotypes of three consecutive passages were assessed.

For miRNA transfection experiments, *mir-290−/−;mir-302~367−/−* iPSC-like cells were plated at 1000 cells per well in 96-well plates in feeder-free media and transfected the next day with 25nM miR-302b mimic and Dharmafect1 transfection reagent per the manufacturer’s protocol (GE/Dharmacon). Cells were passaged every 3-4 days and transfection was repeated one day after passaging.

### Crystal violet proliferation assay

iPSC-like cell lines (*mir-290−/−;mir-302~367+/+* or *mir-290−/−;mir3-2~367−/−*) were plated at 10,000 cells in each well of a 24-well dish. Cells were stained with crystal violet at 24, 48, and 72 hours thereafter. In brief, wells were washed twice with PBS, after which crystal violet solution (0.2% crystal violet (w/v), 2% EtOH in water) was added and incubated at room temperature for 10 min. Wells were rinsed twice in water and incubated in 1% SDS in water until well color was uniform. Level of absorbance of crystal violet as an indication of amount of cellular DNA in each well was read at 570nm on a spectrophotometer (SpectraMax M5) and normalized to a blank control. Data were analyzed using SoftMax Pro software.

### qRT-PCR

Total RNA was collected using TRIzol according to the manufacturer’s protocol. For cDNA synthesis, RNA was treated with DNaseI (Invitrogen) and reverse-transcribed with oligo-dT primers using the SuperScript III kit (Invitrogen). Total cDNA was diluted 1:5, and qPCR was performed using gene-specific primer sets (listed in Supplemental Table 1) and SensiFast SYBR Hi-ROX master mix (Bioline). MiRNA qRT-PCR was performed with the polyA and SYBR Green method as previously described using miRNA-specific forward primers and a 3’ RACE adaptor reverse primer^62^. Primer specificity was verified through analysis of dissociation curves in experimental, no RT, and water-only samples.

### Immunohistochemistry

Cells were fixed for 15min in 4% PFA, washed in PBST (PBS + 0.1% Triton X-100), incubated for 1hr at room temperature with blocking buffer (PBST, 1% goat serum, 2% BSA), then incubated overnight at 4C with 1:500 Nanog antibody (CST8785) in blocking buffer. Cells were then washed in PBST, incubated for 1hr at room temperature in 1:500 secondary antibody in blocking buffer (AlexaFluor 594 goat anti rabbit IgG), washed in PBST with 1:10,000 Hoechst 33342 (Invitrogen), and stored in PBS before imaging using the INCell Analyzer 2000 (GE).

### Genotyping

DNA was extracted by incubating cells in DNA lysis buffer (100mM Tris pH 8.0, 5mM EDTA, 0.2% SDS, 200mM NaCl, 100ug/ml proteinase K) overnight at 55C, precipitating with equal volume isopropanol, washing with 70% EtOH, air-drying, and resuspending in water. PCR was performed using the KAPA HotStart PCR kit (KAPA Biosystems). For *miR-302* Het and KO cells, the 302 wt/mut genotyping F, 302 wt genotyping R, and 302 mut genotyping R primers were used. For *miR-290* WT, Het, and KO cells, the 290 wt/mut genotyping F, 290 wt genotyping R, and 290 mut genotyping R primers were used. For *miR-302~367* CRISPR cells, the 302 CRISPR geno F and 302 CRISPR geno R primers were used. Primer sequences are listed in Supplemental Table 1.

### RNA sequencing and analysis

Total RNA was extracted and purified from cells using TRIzol followed by isopropanol and ethanol precipitation. RNA-seq libraries were generated using the QuantSeq 3’ mRNA-Seq Library Prep Kit FWD for Illumina (Lexogen, CAT#A01172) according to their protocol using 200ng of total RNA for input. We utilized the PCR Add-on kit for Illumina (Lexogen, CAT#M02096) to determine an appropriate number of PCR cycles to amplify libraries. Amplified libraries were quantified using Agilent Tapestation 4200. Libraries were pooled and sequenced using a HiSeq 4000 to obtain single-end 50bp reads. At least 20 million mapped reads per sample were obtained.

Fastq files for all RNA sequencing samples were processed using Nextflow and the nf-core RNAseq pipeline v3.2. In brief, TrimGalore was used on raw fastq files for adapter trimming and quality filtering, samples were aligned using STAR to UCSC mm10 and a gene count matrix was generated using Salmon. Using custom R analysis scripts, the gene count matrix was normalized to counts per million (CPM), and all genes were filtered for genes >1 CPM in at least 6 samples. Differential expression analyses were performed using DESeq2 contrasting all samples for both colonies of a given genotype (4 samples total) to WT. Gene ontology analyses were conducted using Clusterprofiler. To generate cumulative distribution function plots on predicted miR-290 targets, we used all targets predicted in TargetScanMouse for broadly conserved family miR-291-3p/294-3p/295-3p/302-3p containing at least one conserved seed sequence. In preparation for UMAP analysis, removeBatchEffect from Limma was used to batch correct between our RNA-seq data and DGCR8KO data.

## Acknowledgements

We would like to thank Daniel Saw for technical support. This work was supported by funds to R.B. from the National Institutes of Health (R01 GM101180). J.Y. was supported by the National Institutes of Health (predoctoral fellowship F30HD084120) and the UCSF Discovery Fellows fellowship. R.M.B. was supported by the National Institutes of Health (predoctoral fellowship T32 HD007470), UCSF Discovery Fellows fellowship, and ARCS achievement award. R.J.P. was supported by the National Institutes of Health (postdoctoral fellowships T32 HD 007263-27 and F32 HD 070572).

## Data Availability Statement

The raw data that support the findings of this study are available from the corresponding author upon reasonable request. Scripts used for analysis of RNA-seq data are available online through both Zenodo (DOI:<u>10.5281/zenodo.11302307</u>) and GitHub (github.com/ryanmboileau/Ye_Boileau_2024). RNA-seq data generated in this study is available online in the GEO database using the identifier GSE268461. DGCR8KO ESC and matching WT data was acquired from the GEO database using the identifier GSE112767.

## Author Contributions

- Julia Ye: Conception and design, collection and assembly of data, data analysis and interpretation, financial support, manuscript writing
- Ryan M. Boileau: Conception and design, collection and assembly of data, data analysis and interpretation, financial support, manuscript writing
- Ronald J. Parchem: Conception and design, provision of study material, collection and assembly of data, data analysis and interpretation, financial support, manuscript writing
- Robert L. Judson: Conception and design, collection and assembly of data, data analysis and interpretation, manuscript writing
- Robert Blelloch: Conception and design, financial support, provision of study material, data analysis and interpretation, manuscript writing, final approval of manuscript

## Disclaimers

None.

## Grants Acknowledgements

- Robert Blelloch: RO1 GM101180
- Julia Ye: F30HD084120, UCSF Discovery Fellows fellowship
- Ryan Boileau: NIH T32 HD007470, UCSF Discovery Fellows fellowship, ARCS achievement award
- Ronald J. Parchem: T32 HD 007263-27, F32 HD 070572

## Supplemental Figures

**Supplemental Figure 1.**
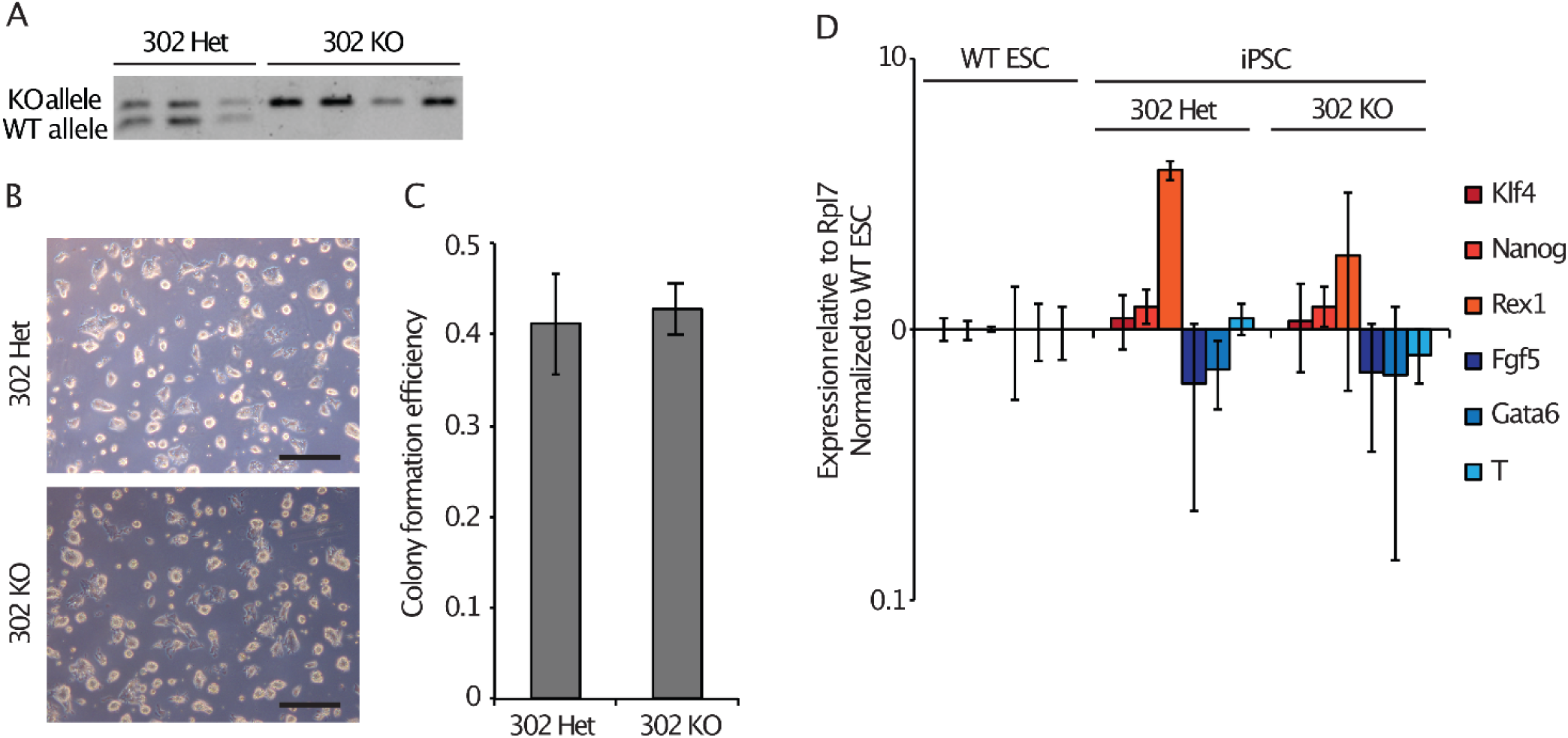
(A) DNA genotyping results for the *mir-302~367* locus in *mir-302* Het and *mir-302* KO iPSC-like cell lines. (B) Representative brightfield images of reprogrammed and expanded *mir-302* Het and *mir-302* KO iPSC-like cells taken at 5x magnification (scale bar represents 500 um). (C) Colony formation efficiency (fraction of cells plated that form colonies) of *mir-302* Het and *mir-302* KO iPSC-like cells. (D) qRT-PCR analysis of naïve pluripotency markers (Klf4, Nanog, Rex1) and primed pluripotency markers (Fgf5, Gata6, T) in WT ESCs and *mir-302* Het and *mir-302* KO iPSC-like cells at P.12. Error bars represent SD of 3-4 biological replicates.

**Supplemental Figure 2.**
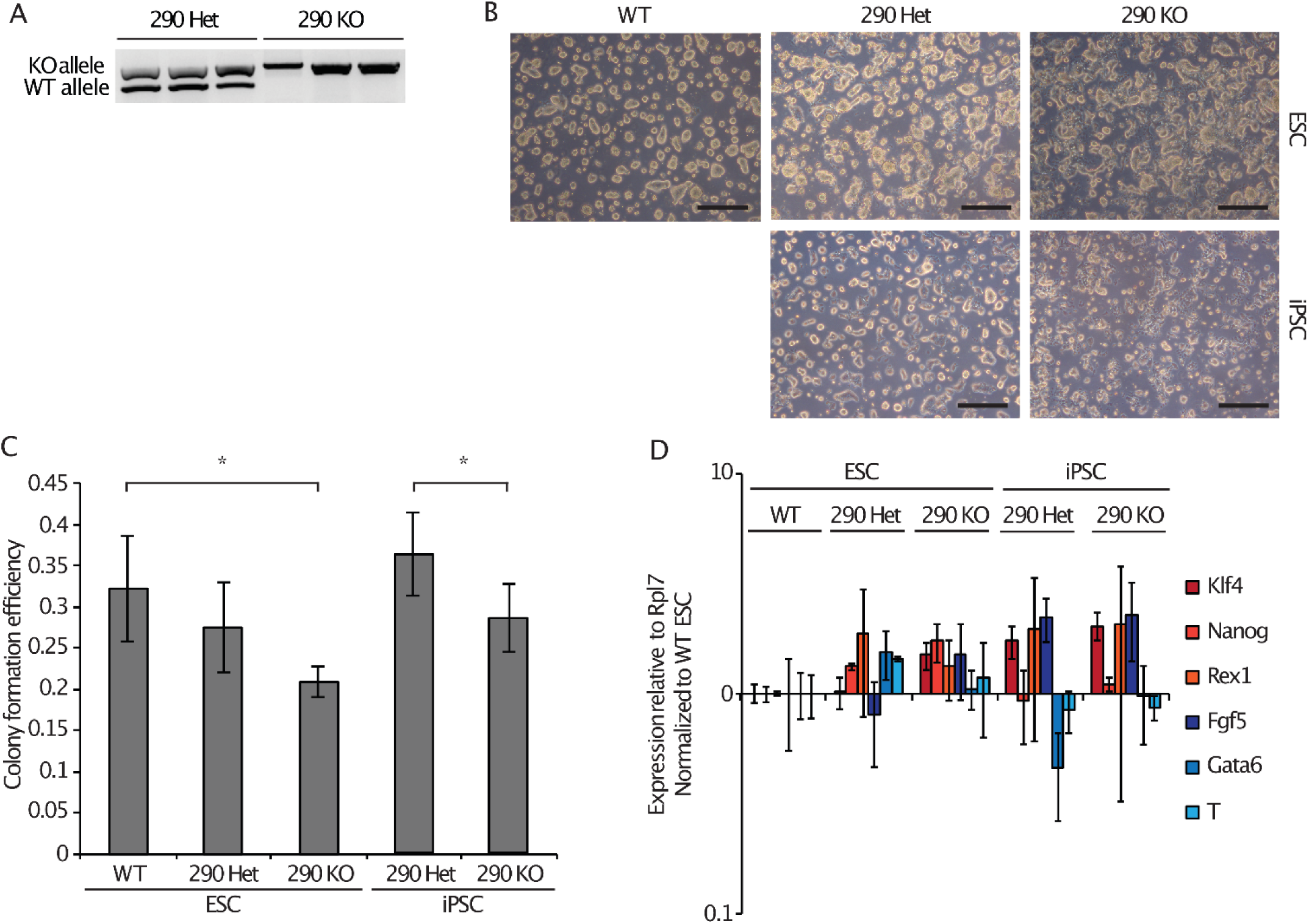
(A) DNA genotyping results for the *mir-290~295* locus in the *mir-290* Het and *mir-290* KO iPSC-like cell lines. (B) Representative brightfield images of reprogrammed and expanded WT, *mir-290* Het, and *mir-290* KO ESCs and iPSCs taken at 5x magnification (scale bar represents 500 um). (C) Colony formation efficiency (fraction of cells plated that form colonies) of *mir-290* Het and *mir-290* KO ESCs and iPSC-like cells. (D) qRT-PCR analysis of naïve pluripotency markers (Klf4, Nanog, Rex1) and primed pluripotency markers (Fgf5, Gata6, T) in WT ESCs and *mir-290* Het and *mir-290* KO iPSC-like cells at P.12. Error bars represent SD of 3 biological replicates. (*) P < 0.05, two-sided t-test.

**Supplemental Figure 3.**
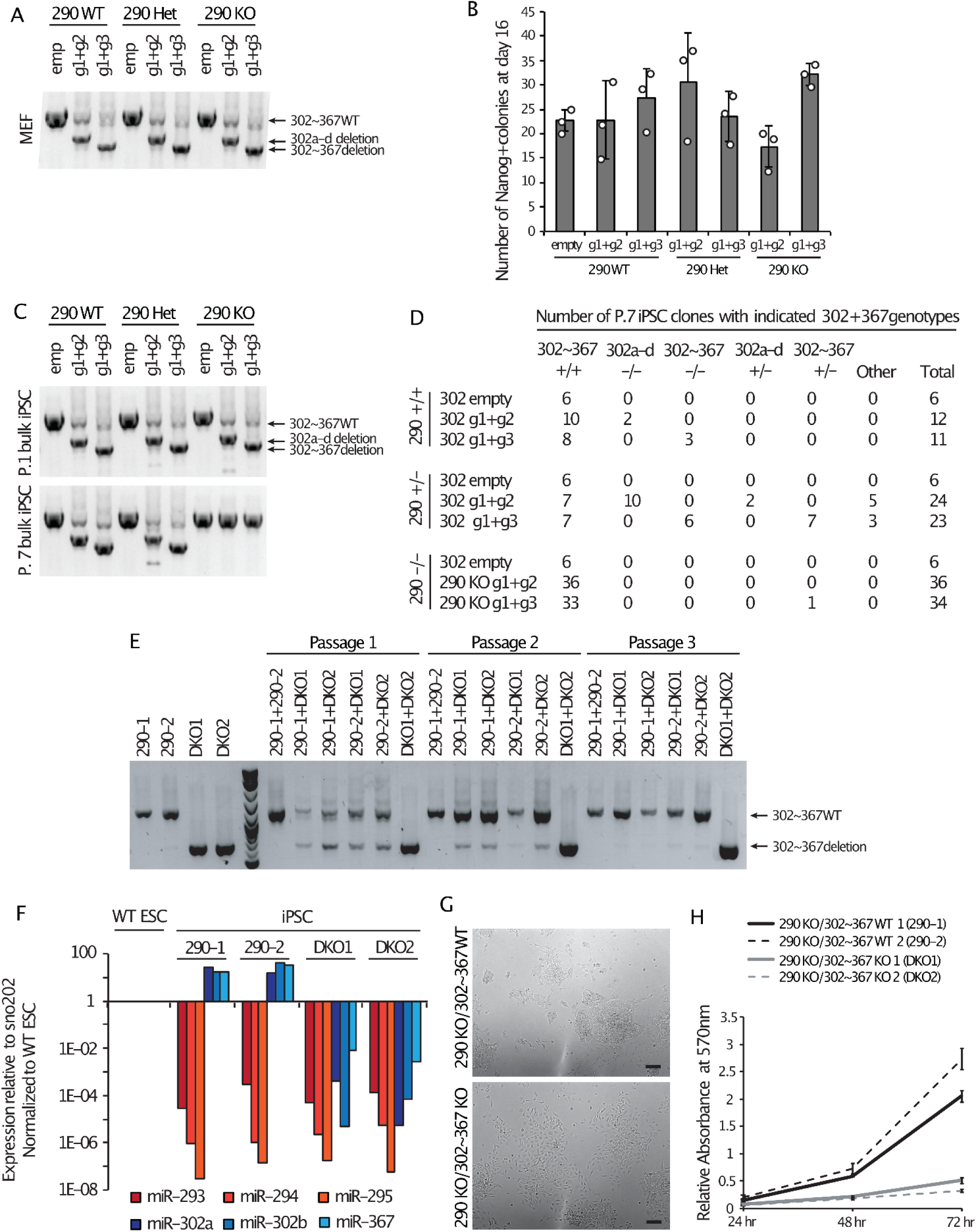
(A) Genotyping results for the *mir-302~367* locus in the starting WT, *mir-290* Het, or *mir-290* KO MEF population. Arrows indicate PCR products representing WT *mir-302~367*, *mir-302a-d* deletion, and *mir-302~367* deletion. (B) Average number of Nanog+ colonies counted per well 16 days after OSK transduction. Cells electroporated with an empty CRISPR construct (“empty”) were used as a control. Error bars represent SD of 3 biological replicates. Circles indicate individual data points. (C) Genotyping results for the *mir-302~367* locus in iPSCs harvested in bulk at P.1 and at P.7. (D) Number of individual iPSC clones of each indicated genotype picked at P.7. “Other” indicates non-WT PCR bands that are not consistent with either *mir-302a-d* or *mir-302~367* deletion. € Representative genotyping results over 3 consecutive passages when equal numbers of *mir-290−/−;mir-302~367+/+* (290-1, 290-2) and *miR-290−/−;mir-302~367−/−* (DKO1, DKO2) iPSCs were initially mixed. (F) Transcript levels relative to internal control and normalized to WT ESCs of mature *mir-290~295* and *mir-302~367* cluster miRNAs in WT ESCs, *mir-290−/−;mir-302~367+/+* (290-1, 290-2), and *miR-290−/−;mir-302~367−/−* (DKO1, DKO2) iPSC-like lines at P.12. (G) Representative images of *mir-290−/−;mir-302~367+/+* and *mir-290−/−;mir302~367−/−* iPSC-like lines taken at 10x magnification (scale bar represents 100 um). (H) Crystal violet absorbance levels in proliferation assay of two *mir-290−/−;mir-302~367+/+* (290-1, 290-2) and two *mir-290−/−;mir-302~367−/−* (DKO1, DKO2) iPSC lines. Absorbance is relative to blanked control. Error bars represent SD of 5 technical replicates.

**Supplemental Figure 4.**
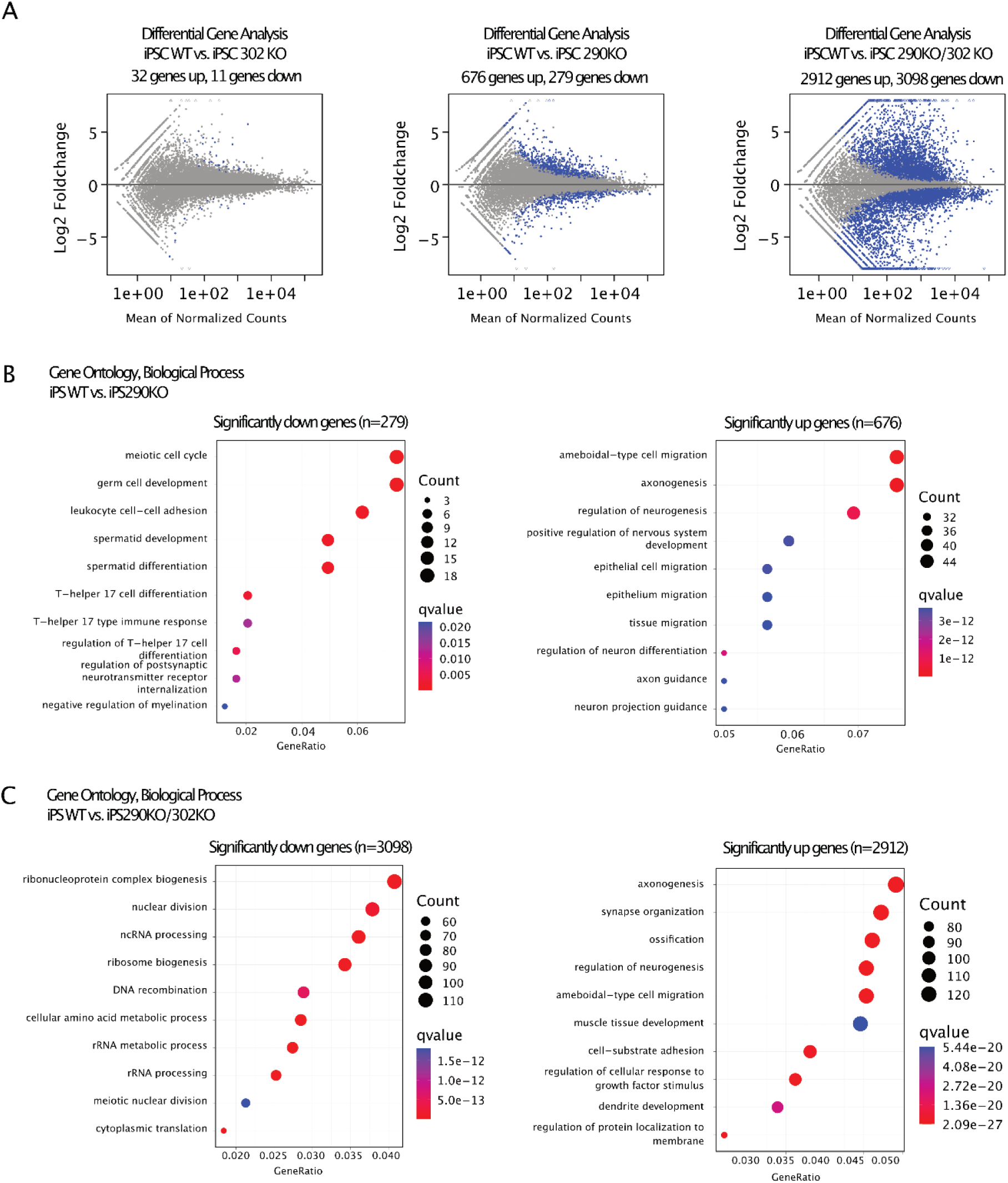
(A) MA plots showing gene expression compared between WT iPSC and either 302KO iPSC-like cells, 290KO iPSC-like cells, or double KO iPSC-like cells. Significantly different genes highlighted in blue (Log2 Foldchange >1, p.value < 0.05 after correction using Benjamini-Hochberg method). (B) Top 10 gene ontology categories enriched using Clusterprofiler on differential genes identified using DESeq2 comparing WT vs. iPSC 290KO. (C) Same as (B) but comparing WT iPSCs vs double KO cells.

**Supplemental Figure 5.**
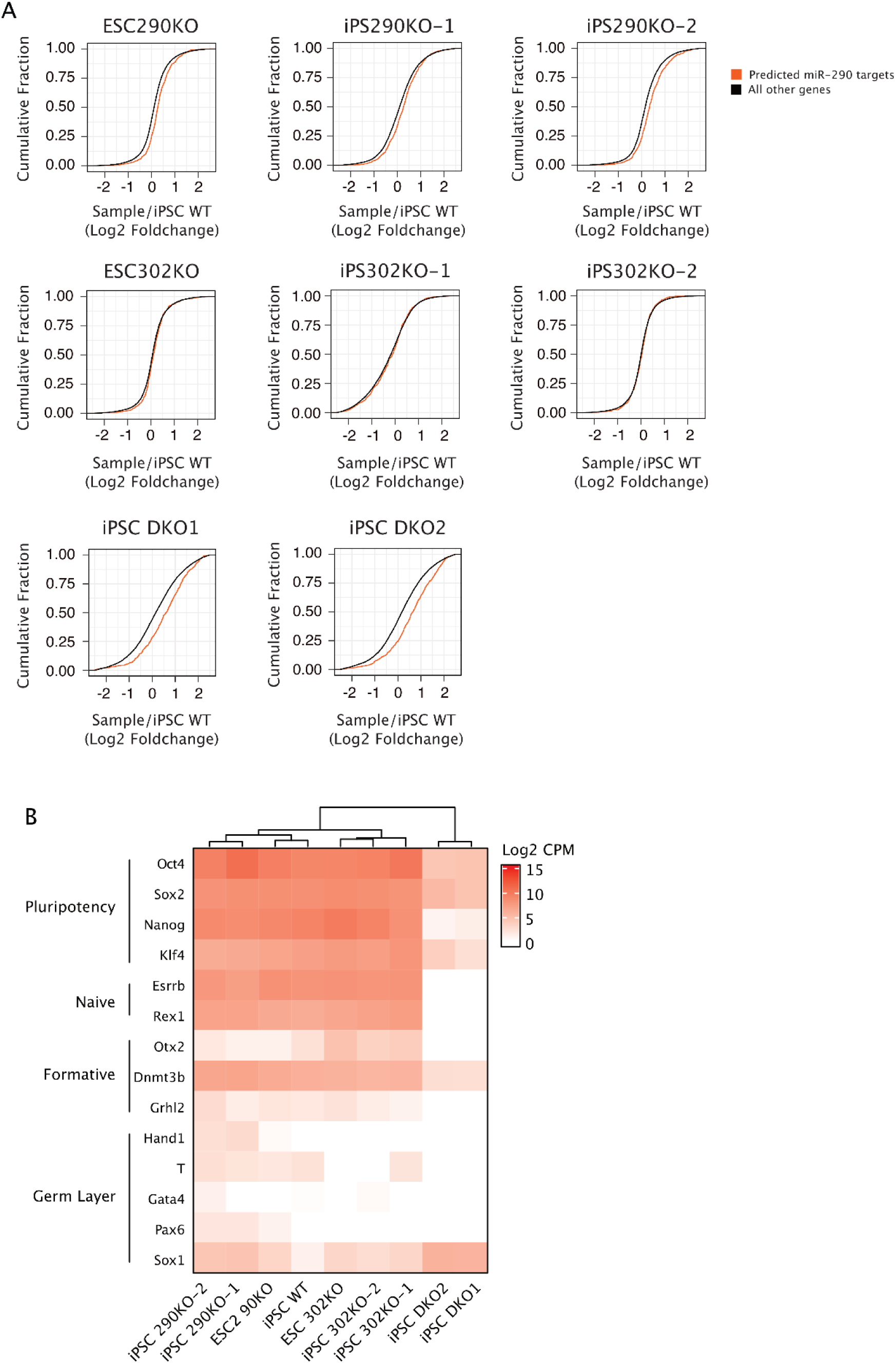
(A) Plots comparing cumulative distribution functions of predicted miR-290/302 family targets (orange) to all other genes detected (black). Values on the X-axis are Log2 Foldchanges for each sample relative to iPSC WT cells. (B) A heatmap showing the absolute level of gene expression for select genes present in Figure 3H. Values represent Log2 counts per million (CPM). The plot is hierarchically clustered by sample.

